# LAT encodes T cell activation pathway balance

**DOI:** 10.1101/2024.08.26.609683

**Authors:** Adam J. Rubin, Tyler T. Dao, Amelia V. Schueppert, Aviv Regev, Alex K. Shalek

## Abstract

Immune cells transduce environmental stimuli into responses essential for host health via complex signaling cascades. T cells, in particular, leverage their unique T cell receptors (TCRs) to detect specific Human Leukocyte Antigen (HLA)-presented peptides. TCR activation is then relayed via linker for activation of T cells (LAT), a TCR-proximal disordered adapter protein, which organizes protein partners and mediates the propagation of signals down diverse pathways including NFAT and AP-1. Here, we studied how balanced downstream pathway activation is encoded in the amino acid sequence of LAT. To comprehensively profile the sequence-function relationship of LAT, we developed a pooled, single-cell, high-content screening approach in which a large series of mutants in the LAT protein were analyzed to characterize their effects on T cell activation. Measuring epigenetic, transcriptomic, and cell surface protein dynamics of single cells harboring distinct LAT mutants, we found functional regions spanning over 40% of the LAT amino acid sequence. Conserved sequence motifs for protein interactions along with charge distribution are critical sequence features, and contribute to interpretation of human genetic variation in LAT. While mutant defect severity spans from moderate to complete loss of function, nearly all defective mutants, irrespective of their position in LAT, confer balanced defects across all downstream pathways. To understand the molecular basis for this observation, we performed proximal protein labeling which demonstrated that disruption of LAT interaction with a single partner protein indirectly disrupts other partner interactions, likely through the dual roles of these proteins as effectors of downstream pathways and bridging factors between LAT molecules. Overall, we report widely distributed functional regions throughout a disordered adapter and a precise physical organization of LAT and interacting molecules which constrains signaling outputs. More broadly, we describe an approach for interrogating sequence-function relationships for proteins with complex activities across regulatory layers of the cell.

## Introduction

In T cell receptor (TCR)-mediated T cell activation, a Human Leukocyte Antigen (HLA)-presented peptide binding event is converted into a complex response involving cytoplasmic signaling pathways, widespread chromatin remodeling, transcription, and the expression, trafficking and secretion of effector proteins^1,2^. The engaged TCR recruits the kinases LCK and ZAP70, leading to phosphorylation and activation of the transmembrane protein linker for activation of T cells (LAT)^2,3^. By rapidly organizing a collection of protein interactors, the extended disordered cytoplasmic tail of LAT relays the single channel signal of peptide-HLA sensing to activation of diverse downstream pathways, including NFAT and AP-1 (**Figure 1A**) ^4,5^. Comparisons of LAT to other signaling proteins have helped identify critical sites of tyrosine phosphorylation and residues required for localization to the plasma membrane; other functional regions, meanwhile, have been nominated through careful consideration of known protein interaction determinants and potential sites of post-translational modification^6–9^. The extent to which other regions of the LAT protein encode function, and the relative contributions of various sites to the activation of different downstream pathways, are not known.

**Figure 1:**
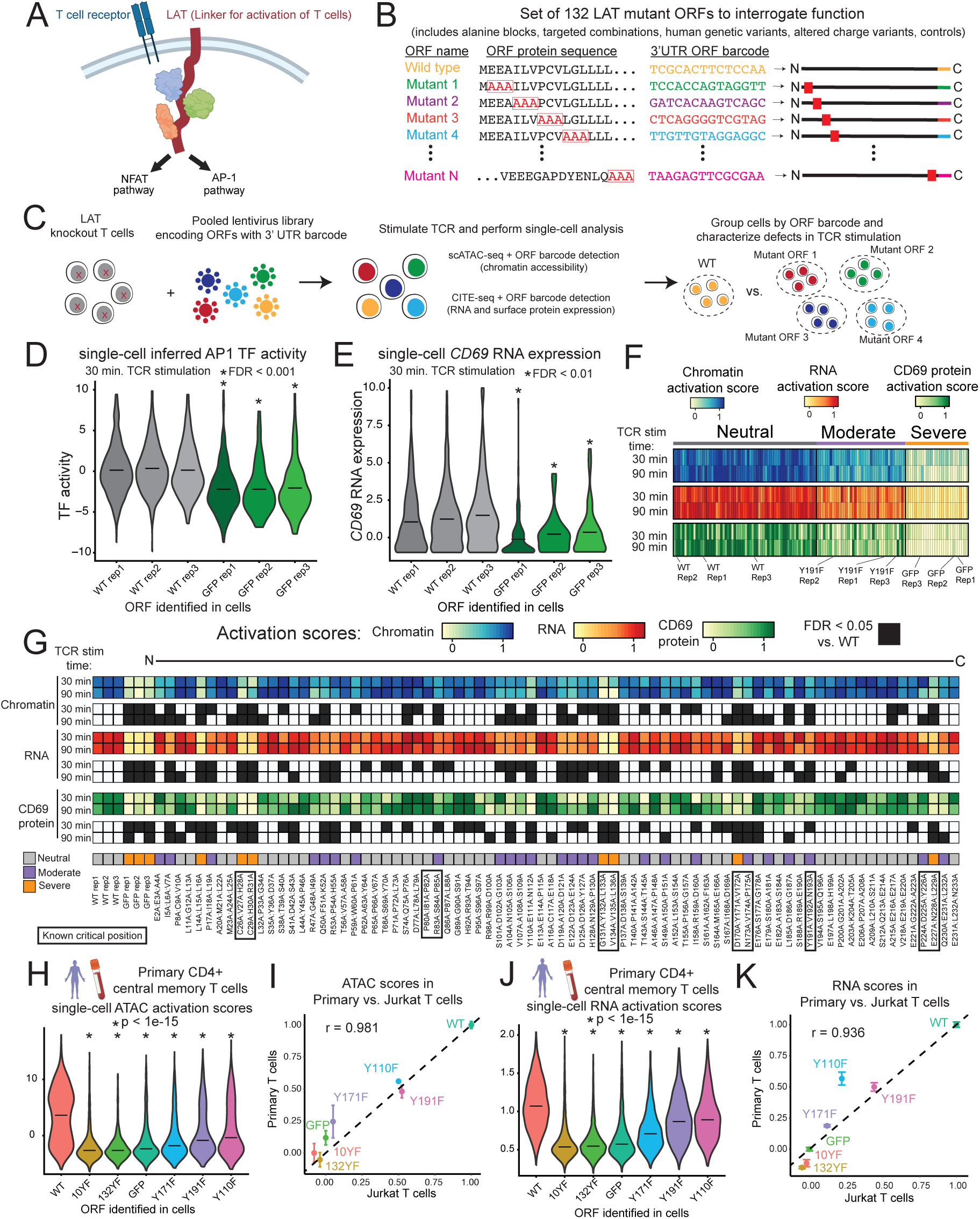
A pooled screen links protein sequence to high-content readouts of parallel LAT activities. (**A**) LAT is a largely disordered membrane-integrated adapter protein which, upon TCR stimulation, aggregates numerous protein interactors to trigger intracellular signaling pathways. (**B**) Triple alanine block mutant ORFs were designed to cover the entire length of LAT. Other ORFs in the library include single and multisite mutants reported in previous studies, variants observed in humans, ORFs altering net charge, and controls. Each ORF is encoded in a cDNA expression construct with an ORF-identifying barcode in the 3’ UTR. (**C**) A single pool of lentivirus corresponding to all 132 ORFs was used to transduce LAT-knockout Jurkat T cells for subsequent TCR stimulation and single-cell epigenomic, transcriptomic, and protein characterization. (**D**) Violin plot and mean inferred AP-1 transcription factor activity from chromatin accessibility for cells assigned to one of the wild type (WT) or GFP replicate ORF barcodes at 30 minutes of TCR stimulation. Replicate ORFs are identical except for the barcode. False discovery rate (FDR) from cell sampling test (See **Methods**). (**E**) Similar to (D), for RNA gene expression of CD69 from the 30 minute TCR stimulation CITE-seq experiment. (**F**) Heatmap of ORF activation scores (mean scores across cells), clustered by k-means. Scores are scaled such that 0 represents the mean of GFP-expressing cells and 1 represents the mean of WT LAT-expressing cells. (**G**) Heatmaps representing chromatin, RNA, and CD69 protein activation scores for ORFs encoding alanine blocks. Scores with an FDR < 0.05 are shaded black in corresponding heatmap rows, and known critical positions from the literature are boxed. Neutral, moderate, and severe indicate cluster labels. (**H**) Violin plot of mean chromatin activation score for primary human CD4+ central memory T cells assigned to each ORF. p-value from KS test, r indicates Pearson correlation. (**I**) Scatter plot comparing chromatin activation scores between Jurkat T cell and primary human T cell models. Error bars represent standard deviation. (**J**) Similar to (H) for RNA activation score. (**K**) Similar to (I) for RNA activation score.

Disordered protein regions, such as the LAT cytoplasmic domain, compose roughly half of the proteome and commonly mediate signal branching^10,11^. These regions can populate an ensemble of conformations, influenced by promiscuous interactions with structured proteins through numerous short linear motifs (SLiMs)^12^. While sequence features such as charge and hydrophobicity are associated with disorder, these regions lack the consistent secondary and tertiary structure of globular domains needed to facilitate functional assignment through homology modeling. While this makes it challenging to predict sequence-function relationships^10,13^, these same features enable disordered regions, including those in LAT, to seed intricate higher-order assemblies that provide spatial and temporal regulation of interacting effector molecules^14^.

Experimental mapping of the relationship between protein sequence and function becomes increasingly challenging when an expected function requires a complex collection of molecular activities across regulatory layers^15^. To date, systematic exploration of protein sequence-function relationships has mostly focused on mutation scanning experiments in which functional output is compared for a library of mutants spanning the length of the protein. This approach has been typically limited to activities directly linked to narrow, easily tracked cellular phenotypes, such as growth or abundance of a single molecular feature, which facilitate the selection of a population of cells^16–20^. Alternatively, bulk genomic measurements can be used to obtain more holistic phenotypic descriptors; however, throughput, cost, and labor limitations associated with these approaches dramatically reduce the scale of mutants that can be queried to a handful of sequence variants selected with strong hypotheses^21^.

Pooled, perturbation-indexed single-cell assays can bridge the gap between traditional genetic screens and low-throughput, information-rich assays^22–24^. Cells genetically modified in a large pool can be analyzed to link perturbations to their effects on complex molecular phenotypes and single-cell heterogeneity. Perturb-seq^22^ is a method to link genetic variants to transcriptomic phenotypes. Similar in experimental design, Spear-ATAC^25^ measures global chromatin accessibility along with a genetic variant barcode in cells enabling associations between genetic perturbations and inferred transcription factor activity. These and similar methods have been employed to explore gene disruption, gene activation, and genetic variation at the epigenomic, transcriptomic, and protein level^25–29^. We sought to extend this general approach to the problem of systematic protein sequence-function mapping.

Here, we developed a pooled screening workflow to assign each region of LAT to its function in organizing aspects of the complex T cell activation phenotype. As LAT represents the branch point of T cell activation signaling, whereby multiple cytoplasmic pathways are controlled by the organization of the LAT signalosome, no single molecular feature encompasses the activity of LAT. To address this, we measured epigenome, transcriptome, and surface protein dynamics in single cells expressing various LAT mutants, enabling us to determine the extent of functional sequence in LAT and how downstream pathways are balanced in cells undergoing activation.

## Results

### A pooled screen links protein sequence to high-content readouts of parallel LAT activities

We designed a library of open reading frames (ORFs) to interrogate the sequence-function relationship of LAT (**Figure 1B**). These ORFs consisted of a mutation scan in which sequential sets of three amino acids were mutated to alanine, as well as a series of negative controls expressing green fluorescent protein (GFP), individual amino acid mutants suggested by previous studies and combinations thereof, and missense mutations observed in humans (**Supplementary Table 1**). We engineered cDNAs encoding each ORF, along with an ORF-specific 14 base-pair barcode with adjacent Perturb-seq^26^ and Spear-ATAC^25^ primer binding sites in the 3’ untranslated region (UTR), in a lentiviral expression vector (**Figure S1A, Supplementary Table 2**). We generated lentivirus in an arrayed fashion from each construct and independently tittered and pooled it for equal representation (**Figure 1C**, **Methods**).

To evaluate the ability of these ORFs to restore LAT function in T cell activation, we first used CRISPR-Cas9 to knock out (KO) LAT in Jurkat T cells and isolated a clone with dual-allele frameshift indels (**Figure S1B, Methods**). Upon re-introduction of wild type LAT, these cells exhibited the expected chromatin and gene expression responses to TCR stimulation, similar to those observed in primary human T cells (**Figures S1C,D**). We next transduced the Jurkat LAT KO cell line with the 132 ORF library lentiviral pool, including the WT, selected for transduced cells, stimulated the TCR with anti-CD3 antibody, and performed single-cell chromatin accessibility (scATAC-seq, with Spear-ATAC^25^ modifications for ORF barcode recovery) or single-cell RNA plus protein (CITE-seq^30^, with Perturb-seq^26^ modifications for ORF barcode recovery) profiling using the 10x Genomics platform. CITE-seq enabled detection of surface expression of the protein CD69, a canonical marker of T cell activation. We analyzed cells stimulated for 30 or 90 minutes to capture early events associated with TCR-proximal signaling (**Figure S1E**). Analysis of the ORF barcodes indicated that individual cells generally express a single dominant ORF (**Figure S1F**). After filtering for cells with high quality scATAC and scRNA features and confident single ORF barcode identification, we retained 20,558 cells for the 30-minute stimulation and 25,734 cells for the 90-minute stimulation in the scATAC experiment and 20,274 and 22,202 cells in the matched CITE-seq experiments (**Figure S1G**). This corresponded to a mean of 152 to 193 cells per ORF in each experiment (**Figure S1H**), substantially exceeding the ∼50 cells per perturbation that previous power analyses suggested are sufficient for robust phenotyping of knockout^22^ or coding variant^26^ effects. Together, these workflows yielded data to link single-cell chromatin accessibility, RNA, and surface protein expression to specific ORF barcodes.

As a first exploration of the ability of these data to inform ORF function, we compared groups of cells expressing WT LAT or GFP as positive and negative controls, respectively. The inferred TF activity of AP-1 (based on chromatin accessibility data) and the expression of the *CD69* RNA and protein, all well-established features gained in T cell activation, were significantly reduced in cells expressing GFP *vs*. WT LAT (**Figures 1D,E, S2A, Supplementary Table 3**)^1,31^. These results indicate that our pooled single-cell ORF screen can successfully distinguish functional and defective restoration of TCR signaling, supporting further interrogation of the 132 ORF LAT variant library.

We next sought to classify ORFs into functional categories. In principle, there are two major ways by which to assess variant effects based on a high-dimensional profile^32^: one uses the full profile or all of its relevant features together as a variant impact score^26^, and the other decomposes it to identify impact of different pathways, genes, or programs^22^. A single global score helps determine if a variant is generally affecting the cell’s phenotype and the overall magnitude of this effect, whereas decomposition helps identify the ways in which this global effect is mediated (including how variants may operate on different downstream pathways). We pursued each approach in turn.

To summarize the high dimensional chromatin and RNA data, we first established a score based on the top differential TFs and genes between cells expressing WT LAT and GFP in TCR stimulation (**Methods**). (We discuss individual features further below.) Briefly, in each such score, in each cell the scores for the top 50 respective features (accessible TF motifs for chromatin, genes for RNA) were averaged, and then further averaged across all the cells with one ORF. These chromatin and RNA scores, as well as CD69 protein levels were then scaled for the cells with each ORF such that 0 represents the mean level in GFP-expressing cells and 1 represents the mean level in WT LAT-expressing cells (**Supplementary Table 4, Methods**). Using these scaled scores, we performed k-means clustering and identified three groups of ORFs (**Figures 1F, S2B**). The first cluster contained all three WT LAT replicates and 61 other ORFs exhibiting generally neutral effects on activation by this score. A second cluster, containing all three replicates of Y191F (mutation of tyrosine at position 191 to phenylalanine, known to disrupt a protein interaction motif ^5,33^ and 37 other ORFs, showed moderate loss of activation across modalities. The most severely defective ORFs fell into a third cluster, which contained all three GFP replicates as well as 25 other ORFs. Together, these data support the reproducibility and extent of defects observed in the screen.

An important consideration when interpreting the impact of mutations on protein functions is that mutations may influence protein stability and ultimately steady state expression levels. This mode of defect could provide a challenge to interpreting the effect of mutations on function through altered molecular activity such as protein interactions or trafficking^15^. To address this possibility, we performed a modified version of the inCITE-seq experiment, which uses a cell permeabilization approach compatible with CITE-seq to enable the detection of protein epitopes within cells via DNA oligo-conjugated antibodies ^34^. We used an anti-FLAG CITE-seq antibody targeting the N-terminus of each ORF in our library to quantify protein expression (**Figure S2C**, **Methods**). Overall protein expression was not related to a combined chromatin and RNA activation score (FDR = 0.337), indicating that our results more likely reflect altered molecular activities of LAT rather than alterations in abundance.

To understand the distribution of functional sequence throughout LAT, we examined the activation scores for each ORF in the mutation scan in linear order (**Figure 1G**). Mutants spanning known critical residues exhibited strong defects across each modality, including: cysteines 26 and 29, which are palmitoylated and required for membrane localization^35^; tyrosine 132, which is phosphorylated and serves as a docking site for PLCG1^5,33^; and tyrosines 171, 191, and 226, which are also phosphorylated and bind other critical adapter proteins including GRB2^2,5,33^. Additional defects included the proline rich stretch of PIPRSP (residues 80-85), which binds the SH3 domain of LCK to promote ZAP70 localization to LAT^8^. These expected defects indicate that our experiment captures known biology and allows for exploration across LAT.

Beyond these known regions, we discovered several others of previously uncharacterized function. Indeed, over 40% of mutant scan blocks fell into the moderate or severe clusters. To validate the function of previously uncharacterized sites, we selected eight positions, separately transduced cells with the corresponding individual mutant ORFs, stimulated them with anti-CD3 antibody, and performed flow cytometry analysis for CD69 surface protein expression (**Figure S2D)**. These mutants all conferred activation defects, thus confirming that our approach can identity novel functional sites within LAT.

To extend our results beyond the Jurkat T cell model, we asked whether similar mutant ORF phenotypes manifested in primary human CD4+ memory T cells. We knocked out endogenous LAT in primary T cells, delivered six LAT variants and GFP, pooled cells, and performed single-cell ATAC-seq and RNA-seq with ORF barcode detection (**Methods**). After distinguishing central and effector memory T cell states using canonical marker genes, we scored each cell for chromatin and RNA activation, and found that LAT variants recapitulated defects at a similar severity observed in the Jurkat T cell screen (**Figures 1H-K**, **S2E**). Together, these results indicate that our high-content screen recovers known functional sites in LAT and discovers novel sites with relevance in primary human T cell activation.

### Determinants of sequence function in LAT

We next sought to leverage this comprehensive set of mutations to understand sequence features underlying function in LAT. We first examined sequence conservation to highlight regions conferring critical functions. We used the ConSurf tool to score each alanine block by mean conservation score and found, as expected, that known critical residues such as cysteines 26 and 29 and tyrosines 132 and 171 resided in highly conserved blocks (**Figure 2A**)^36^. All mutants from the severe cluster were highly conserved (p < 1×10^-3^ in comparison to neutral cluster mutants using a combined chromatin and RNA score) (**Figure 2B, S3A**). While moderate cluster mutants were significantly more conserved than neutral mutants (p < 0.05), both moderate and neutral mutants spanned a broad range of conservation levels, from high to the most poorly conserved positions in LAT. These results indicate that conservation alone is not strictly linked to function in our experiment, raising the possibility that LAT is adaptable in that it contains functional residues with high sequence variation in evolution – such residues may play species-specific roles^37^. Residues with high conservation and neutral scores in our experiment may contribute to LAT activity in other contexts, such as particular stages of T cell differentiation or in NK or mast cells where LAT is required for FcR signaling^38,39^. The presence of highly conserved yet functionally neutral residues (L11:G12:L13) in the transmembrane domain suggests that interactions within the membrane may also be context-specific (**Figure 2C**). Overall, mutants conferring severe defects were tightly conserved, likely representing universal functions of LAT across cell types and species, while moderately defective mutants may play context-specific roles.

**Figure 2:**
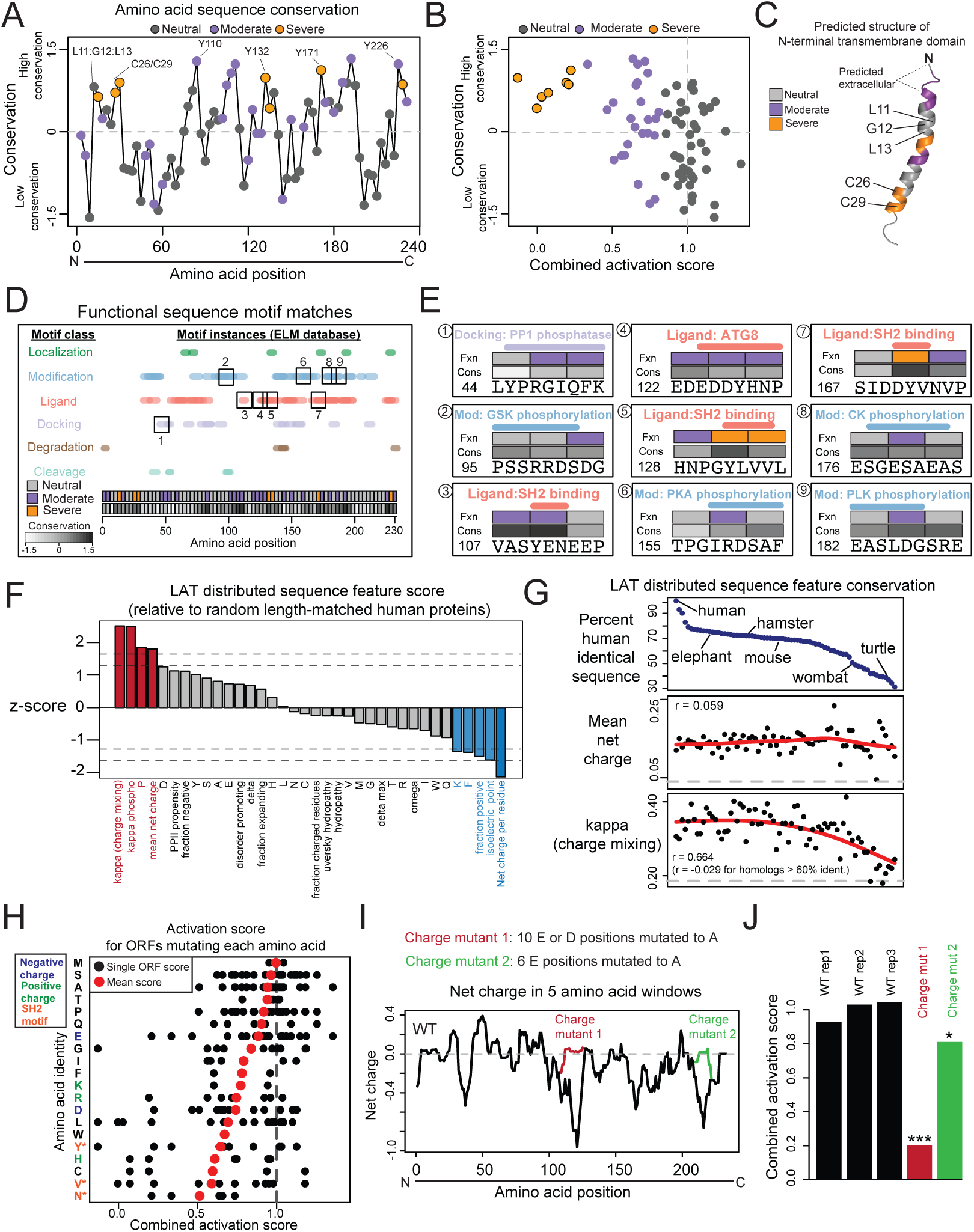
Determinants of sequence function in LAT. (**A**) Amino acid sequence conservation of triple alanine blocks was calculated using the ConSurf tool. ConSurf computes a normalized score for each residue in the protein, whereby the mean value across all residues is zero and the standard deviation is one. Plotted values represent the mean conservation score for residues in the three-residue block. (**B**) Scatter plot of combined activation score (from chromatin and RNA scores) vs. conservation for each alanine block. (**C**) AlphaFold prediction of LAT transmembrane domain structure. (**D**) Motif position matches from the Eukaryotic Linear Motif (ELM) database. Boxed ligand motifs are displayed in detail in (E). (**E**) Detailed examples of ELM motif matches in LAT. The number in the bottom left of each square indicates the starting amino acid position in LAT. Fxn refers to the functional categories of Neutral, Moderate, or Severe. Cons refers to conservation score as displayed in (D). (**F**) Bar plot indicating the z-score of various sequence features for LAT compared to 100 random length-matched human proteins. Feature values were computed by localCIDER (Materials and Methods). Vertical lines at 5 and 10 percentiles. (**G**) Scatter plots of CIDER distributed amino acid sequence features for LAT homologs, ordered by amino acid sequence identity with human LAT. Pearson correlation (r) calculated between sequence identity percent and each feature. Red line indicates a smooth spline calculated in R and gray line represents the mean of random length-matched proteins. (**H**) For each amino acid, dots represent each ORF in which that amino acid is mutated at least once, located on the x axis position indicated the combined activation score for that ORF. Red dot indicates the mean across all ORFs mutating a particular amino acid. Asterisk indicates FDR < 0.05 compared to mean of score for all residues based on permuting residue positions. (**I**) Running charge (mean within five amino acid windows) for WT LAT and charge mutants. (**J**) Combined activation scores for each charge mutant. (***, FDR < 0.05 in at least one time point of chromatin, RNA, and CD69 protein samples; *, FDR < 0.05 in one time point of CD69 protein sample, **Supplementary Data 4**).

Conserved sites in disordered proteins often harbor short motifs controlling interaction with other proteins, post-translational modification, or sub-cellular trafficking^11^. To characterize these regions, we used the Eukaryotic Linear Motif database which identified motifs throughout LAT, mostly in the ligand-binding and post-translational modification categories (**Figure 2D**, **Supplementary Table 5**)^12^. Nearly all functional regions overlapped at least one motif, and this analysis again identified an overlap between tyrosine-containing SH2 domain-binding motifs and sites that have severe phenotypes and are highly conserved (**Figure 2E**). In addition to tyrosine motifs, we found motif matches potentially explaining the function of several novel sites that are moderately conserved and associated with moderate phenotypes, involving phosphorylation addition/removal and protein interaction.

Beyond short sequence motifs, disordered proteins have been shown to exhibit conserved biophysical features determined by their total sequence, which are critical for function and may be independent of the conservation of local sequence^40,41^. To nominate such features with potential roles in LAT, we used the CIDER tool (**Methods**)^42^. Several features were significantly distinct from random length-matched human proteins, such as charge mixing, mean net charge per residue, fraction of positively charged residues, proline content, and phenylalanine content (**Figure 2F**). Further supporting the importance of these features, mean net charge per residue was conserved across homologs, extending to roughly 30% sequence identity, while charge mixing (a measure of blocks of charge) was conserved across homologs down to roughly 60% sequence identity (**Figure 2G**). The high fraction of disorder promoting residues and low fraction of positively charged residues were also conserved (**Figure S3B,C**). Together these measures highlight the extent and distribution of negative charge as important features of LAT, consistent with a previous study of *in vitro* reconstituted signaling^43^.

To understand the functional consequences of amino acid biophysical features in LAT, we calculated the average activation score for alanine blocks mutating each amino acid (**Figure 2H**). Tyrosine (Y), valine (V), and asparagine (N) were the only significantly defective residues (FDR < 0.05 for each), consistent with their repeated occurrence in LAT’s well-characterized SH2 domain binding sites^44^. Mutating individual instances of charged residues, however, did not generally alter LAT function, suggesting that the total extent and distribution of charge may be important. To investigate this possibility, we examined two ORFs designed to drastically alter regions of concentrated negative charge (**Figure 2I**). Charge mutant one (ORF ID 110) converted 10 proximal E or D positions to neutral residues, while charge mutant two (ORF ID 111) similarly converted six residues, resulting in an increased net charge from −28.9 to −18.9 and −22.9, respectively. Charge mutant one resulted in consistent defects (FDR < 0.05 in at least one time point in all three modalities) and was in the severe cluster, while charge mutant two exhibited a significant yet more moderate effect (**Figure 2J**). Consistent with previous findings, these results indicate that the total extent, and likely the linear patterning of negative charge along LAT are critical features^45^.

Together, these results highlight several key functional determinants of LAT. While conservation is a dominant feature of the most critically functional regions involving the transmembrane domain and tyrosine-based SH2 domain binding motifs, more moderately functional regions may exhibit a range of evolutionary or contextual roles and potential molecular mechanisms. Further, distributed biophysical sequence features such as charge patterning support a mechanism independent of local sequence identity which may involve partitioning of LAT-interacting proteins based on charge^43^.

### Functional interpretation of natural human genetic variants

We reasoned that our experiment could also inform the interpretation of human genetic coding variants in the LAT locus. To this end, we included in our library four missense mutations reported in ClinVar, a database of potentially clinically relevant variants^46^. None of these variants had been experimentally evaluated, and their predicted functional impacts from tools summarized in VarSome (including SIFT,

PROVEAN, and PrimateAI-3D) ranged from likely benign to pathogenic^47^. In our data, three of the four ClinVar variants (P59A, P82L, and P141L) conferred modest yet statistically significant differences in activation compared to WT LAT as determined by chromatin accessibility, RNA levels, or CD69 protein expression (**Figure S3D**).

Reasoning that an alanine block serves as a proxy to inform the function of an overlapping missense variant, we extended these findings by leveraging the alanine block scan mutants to examine potential functions of all missense variants in LAT observed at least twice in humans in the gnomAD database (**Figure S3E**)^48^. Notably, the block corresponding to V7F, which was found at the second highest allele frequency (8×10^-4^) and in two individuals as homozygous, was in the moderate defect cluster (combined activation score 0.58, FDR < 0.05 across all modalities and time points).

V134M, which is found at a low frequency (1.99×10^-5^), overlapped a block that conferred a severe activation defect (combined activation score −0.13, FDR < 1×10^-3^ across all modalities and time points), likely through disruption of the Y132-associated SH2 binding motif which recruits PLCG1. Overall, our data help with functional interpretation of human genetic variants and nominate several variants as potentially pathogenic.

### LAT encodes downstream pathway balance

A hallmark of T cell activation is the simultaneous, properly balanced induction of several signaling pathways leading to hundreds of differentially expressed genes. Thus, for example, two LAT variants can achieve similar ‘moderate’ scores either because they have the same partial impact on all induced pathways, or because they each have a severe impact on a different subset of pathways. We thus sought to understand whether individual sites in LAT, associated with particular sequence features as described above, relate to one or more distinct pathways. LAT could trigger downstream pathways in a modular fashion, by which molecular events at distinct regions of LAT are independent and contribute to distinct pathway outputs, or in a coordinated fashion, in which LAT must form a fully functional signalosome to trigger activation of any and all downstream pathways (**Figure 3A**). We hypothesized that analyzing how mutants affect the activities of individual TFs associated with particular pathways would enable mapping LAT sites to pathways and provide support for either a modular or coordinated model (**Figure 3B**). In examining any two TF activities reflecting distinct pathways, cells lacking LAT completely will have a severe defect in both activities. In a modular model, however, mutants may affect one TF activity yet not necessarily affect the other; in the coordinated model, any mutant affecting one TF will correspondingly affect a second TF.

**Figure 3:**
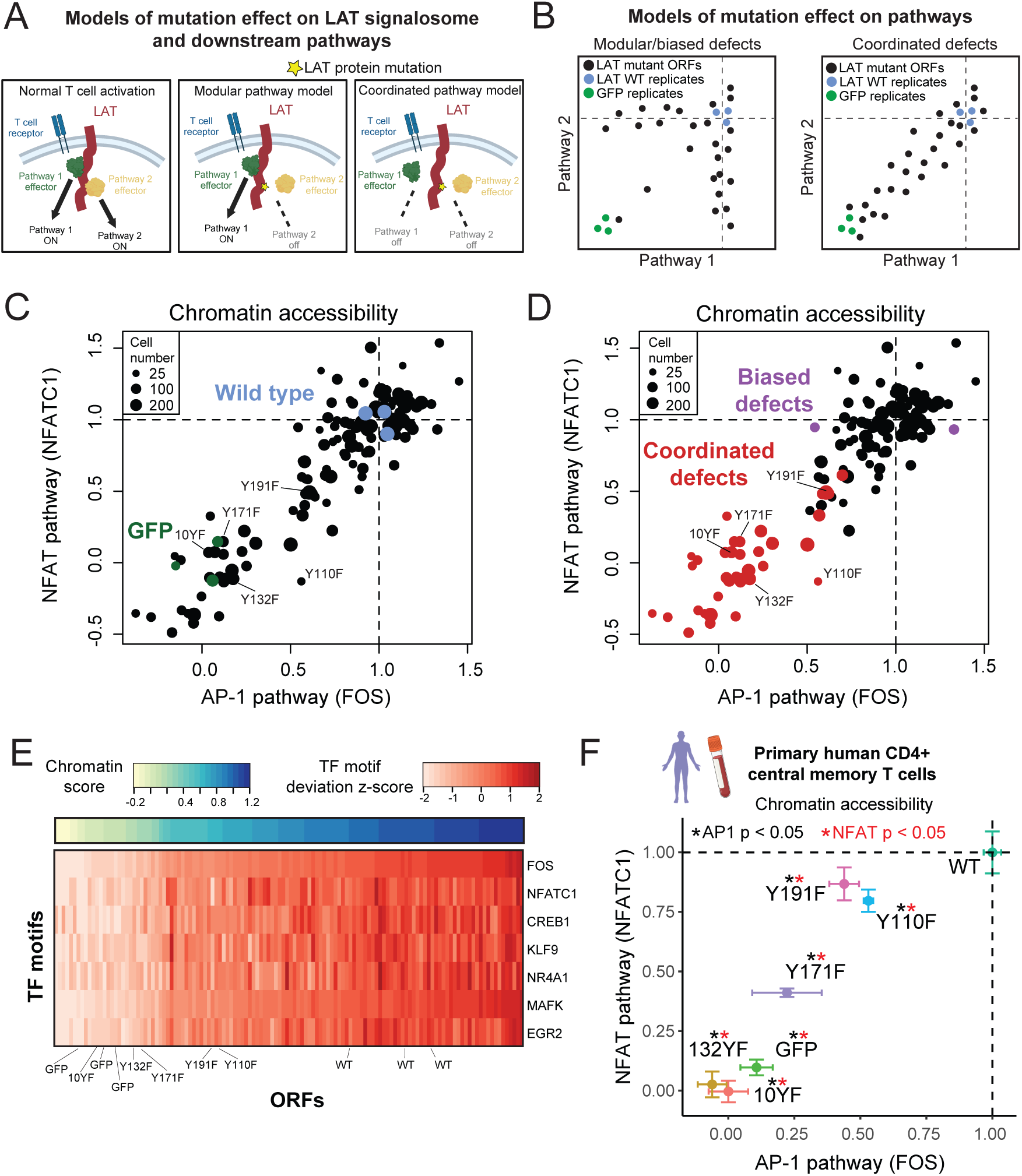
LAT encodes downstream pathway balance. (**A**) Models of LAT with interacting proteins which mediate intracellular signaling pathways controlling chromatin, RNA, or protein features. The complex of LAT with interactors (“signalosome”) could control downstream pathways in a modular or coordinated fashion. In a modular model, mutation in one region of LAT and disruption of a particular interactor will disrupt one downstream pathway while leaving others active. In a coordinated model, mutation in one region of LAT which disrupts a particular interactor may disrupt other interactions or pathway activities (either directly through higher-order physical interactions or indirectly through signal cross-talk). (**B**) Expected results for the models proposed in (A). Modular or coordinated signaling will exhibit distinct patterns of mutant effects on pairs of pathway activities measured in the screen. (**C**) Scatter plot of the accessibility of two chromatin features (inferred TF activity, averaged across cells expressing a particular ORF) representing central pathways of T cell activation. FDR from permutation sampling test. ORFs supported by at least 50 cells are displayed. (**D**) Scatter plot of the same data as in (C), with ORFs labeled as exhibiting balanced or biased defects across AP1 and NFAT pathways. Balanced defects exhibit statistically significant defects in both AP1 and NFAT pathways. (**E**) Heatmap of inferred TF activity for TFs representing motif families that increase in T cell activation. ORFs (columns) are ordered by chromatin activation score. (**F**) Scatter plot of AP1 and NFAT TF activity in primary human CD4+ central memory T cells. Error bars represent standard deviation across replicates, p-value from KS test of single-cell values.

With these models in mind, we compared inferred TF activities from chromatin accessibility data of the AP-1 family member FOS and the NFAT family member NFATC1, representing two of the critical pathways engaged in T cell activation (**Figure 3C**). We observed a linear relationship between the two inferred TF activities, whereby mutants disrupting AP-1 activity consistently disrupted NFAT activity to a similar extent. This was the case not only for severe defects (which can indicate complete loss of function) but also for intermediate, moderate ones, further supporting a coordinated model. Overall, 38 mutants were coordinated (conferring significant comparable defects in both pathways) and only two mutants were biased (conferring a defect in one pathway and not another; **Figure 3D**). Beyond AP-1 and NFAT, the TFs representing other families induced in T cell activation exhibited similar coordinated defects (**Figure 3E**). Further, mutants conferring coordinated defects in the Jurkat T cell screen all had significant defects in both AP-1 and NFAT pathways in primary human CD4+ central memory T cells (**Figure 3F**). Together these results indicate that LAT mutations generally confer coordinated, balanced defects in downstream pathway activities and thus LAT does not exhibit a simple mapping between individual sites and corresponding pathways.

### Indirect disruption of protein interactions underlies balanced defects

Given the extent of broad, coordinated defects from single-site mutations and the potential biological implications of enforcing pathway balance, we sought a molecular explanation of how LAT signalosome activity could be holistically sensitive to any individual defect. We reasoned that defects may be conferred through disruption of direct binding proteins and thus set out to comprehensively map which proteins may directly bind LAT and at which sites. Starting from a list of 10 LAT interacting proteins identified in an affinity purification mass spectrometry dataset, we used AlphaFold-Multimer to predict interaction structures and nominate sites of direct binding^49,50^. Of the 10 proteins, four had AlphaFold-Multimer support as direct binders to segments of the disordered cytoplasmic tail of LAT: PLCG1, GRB2, GADS, and GRAP (**Figure S4A**). Each of these proteins contains phospho-tyrosine binding SH2 domains and have some evidence for binding a subset of LAT tyrosines^6,51^. To determine which LAT tyrosines bind which of these proteins, we reran AlphaFold-Multimer with LAT and the isolated SH2 domains, and scored each predicted structure based on proximity of the SH2 domain to a single LAT tyrosine (**Methods**). While PLCG1 was predicted to bind LAT position 132 exclusively, GRB2, GADS, and GRAP were predicted to permissively bind at any of positions 110, 171, 191, and 226 (**Figures 4A,B**, **S4B-F**).

**Figure 4:**
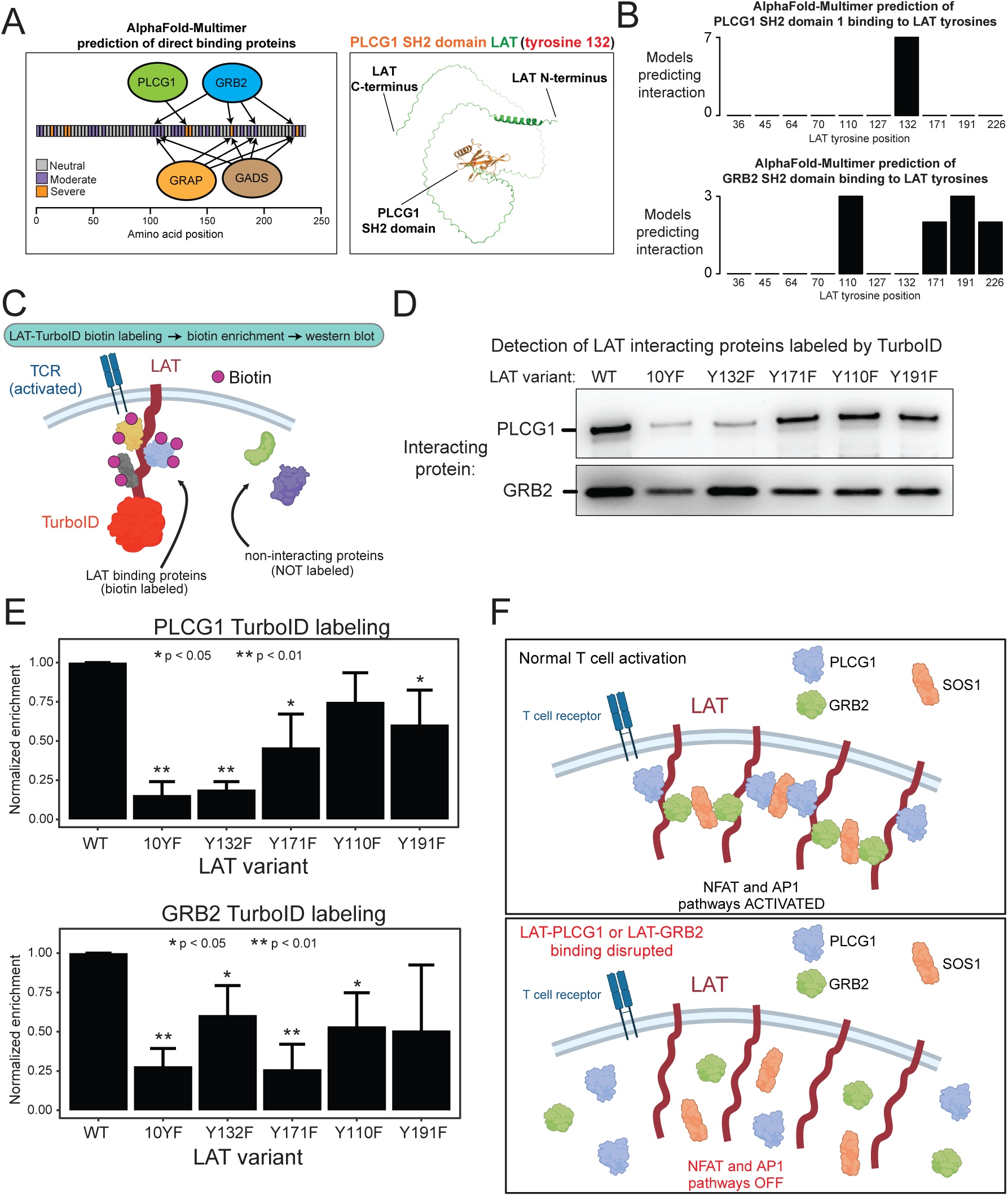
Indirect disruption of protein interactions underlies balanced defects. (**A**) Left: Schematic of binding sites of SH2 domains predicted by AlphaFold Multimer. Right: predicted structure of LAT interacting with the PLCG1 SH2 domain. (**B**) Counts of models (out of 10 total predicted models) for each LAT interaction with an SH2 domain. Models were scored as interaction with a particular LAT tyrosine based on exclusive proximity of 10 angstroms. (**C**) Schematic of LAT-TurboID proximity labeling experiment. LAT fused to TurboID was expressed in Jurkat cells. Cells were activated by pervanadate stimulation for 10 min, and biotinylated proteins were detected by western blot of the streptavidin enriched lysate. **(D)** Representative western blots from four replicate labeling experiments detecting PLCG1 and GRB2 abundance in streptavidin-enriched samples. (**E**) Quantification of band intensity from four replicate labeling experiments. Error bars represent standard deviation and p-values were calculated by t-test. (**D**) Model of LAT interaction with partner proteins. Disruption of one interaction has indirect effects on distinct interactors, resulting in balanced loss of pathway activation.

One possible explanation for multi-pathway, coordinated defects from mutating one site on LAT could be that loss of protein binders from the mutated site also leads to loss of the binding of other proteins from distant sites, for example because signalosome formation is an all-or-none scenario. Even if the proteins binding each site trigger distinct downstream pathways, a defect in one site would be functionally similar to a defect in another site. Given the established role of PLCG1 phospholipase enzymatic activity mediating NFAT pathway activation and GRB2 as an adapter recruiting SOS, the critical guanine nucleotide exchange factor for Ras controlling MAPK and AP-1 signaling, we sought to explore how these proteins, which appear to bind mutually exclusive sites on LAT, may be functionally linked.

We developed a biotin labeling experiment to quantitatively assess LAT interactor proximity in living T cells. Using TurboID fused to the C-terminus of LAT, we reproducibly and selectively detected PLCG1 and GRB2 interaction with LAT in an activation-dependent manner (**Figures 4C**, **S4G-H**). As expected, the LAT 10YF mutant (converting all ten tyrosine residues to phenylalanine) significantly impaired LAT interaction with either PLCG1 or GRB2. Similarly, as expected from previous studies and our AlphaFold modeling, LAT Y132F phenocopied 10YF loss of PLCG1 interaction, indicating that Y132 is the single binding site for PLCG1 (**Figure 4D,E**). Mutations of Y171 and Y191 also confer PLCG1 loss, potentially through disrupting interaction with GRB2 or GADS, which stabilizes the LAT-PLCG1 interaction through the adapter SLP-76^33^. While mutation of expected GRB2 binding sites Y171F and Y110F impaired GRB2 interaction, we found that mutation of Y132, the exclusive PLCG1 binding site, also disrupted GRB2 interaction. Together, our results support a model where mutation of a single site can disrupt protein binding events, and thus their related signaling pathways, at distinct, distant sites on LAT (**Figure 4F**).

## Discussion

Mapping the relationship between protein sequence and function is a fundamental problem in biology. Here, we employed a high-content screen to link the amino acid sequence of the adapter protein LAT to its multifaceted function branching TCR stimulation to numerous downstream pathways. Mutating critical sites of tyrosine phosphorylation and membrane-proximal regions required for cell surface trafficking conferred a near complete loss of activation while mutating a broader collection of sites had more moderate defects.

Individual signaling pathways appear coordinated – LAT mutants affecting one pathway generally affect others to a similar extent. This coordination may occur at the LAT signalosome itself due to the overlapping roles of interacting proteins as pathway-specific effectors (adapters or enzymes) and crosslinking agents bridging LAT molecules to promote cluster formation at the membrane ^43,52^. With SOS as an intermediate or directly, GRB2 and PLCG1 have been shown to promote clustering of phosphorylated LAT, which may provide a temporary environment, protected from CD45 phosphatase activity, in which NFAT/Ca^2+^ and MAPK pathways can be initiated^14,53^. Indeed, this environment may promote dwell time of LAT-interacting proteins at the membrane, a highly sensitive factor regulating SOS and other signaling systems^54,55^.

These coordinated outputs support a model in which LAT recruits and organizes interaction partners in a defined holistic assembly, the activity of which depends on its precise composition (**Figure 4F**). This system may be a mechanism to constrain the types of signaling outputs of T cell receptor stimulation in the face of a wide range of environments. In contrast, a recent study of the Toll-like receptor response found that downstream pathways, while triggered at the same adapter assembly, were controlled in a modular, independent fashion^56^. Future efforts will be required to understand how distinct adaptive and innate immune responses are organized to achieve optimal pathway balance, noise tolerance, and kinetics, among other features encoded in adapter-effector systems.

Our data also likely capture an incomplete picture of LAT sequence-function due to the role of LAT in signaling across various T cell developmental stages and states, as well as in NK and mast cells. While our Jurkat T cell line stimulation system recapitulates the core features of primary human memory T cell activation, and we validated key finding in primary T cells, future experiments in primary T cells of distinct differentiation and memory states, as well as cells experiencing chronic antigen stimulation from infection or cancer, may reveal unique interactions between LAT and TCR proximal signaling machinery that is differentially expressed or regulated in these states. Beyond TCR signals, a collection of co-signaling molecules may shape the proximal signaling environment and shift the sequence-function map to achieve altered pathway balance or activation kinetics. In NK and mast cells, LAT responds to Fc receptor stimulation and may cooperate with distinct adapters and effectors^38,39^.

Our approach may be extended by capturing further molecular and cellular aspects of T cell activation. Improved CITE-seq methods to measure intracellular phosphorylation status and protein-protein interactions within the LAT signalosome would provide more direct mechanistic insights into how LAT regulates and organizes interacting molecules^57,58^. A hallmark of LAT function is formation of short-lived clusters or condensates which could be detected in a pooled screen format using an optical readout^59^. Additional measurement of molecules mediating immune function, such as cytokines TNFα and IL-2 in CD4+ T cells, would shed light on potential post-transcriptional and post-translational regulation.

Future developments increasing the scale of single-cell assays will enable higher resolution queries through single alanine and deep mutational scans, as well as combinatorial mutants to uncover genetic interactions. Mutations may also be achieved via base or prime editing of the endogenous gene locus, although strategies to improve efficiency and detect heterozygotes will be required^60^.

In summary, we describe a sequence-function map of the adapter protein LAT, which is required for T cell activation. By linking LAT sequence regions to high-dimensional readouts across modalities, we captured a complex set of functions associated with this single protein. Defects associated with LAT mutation are generally coordinated across downstream pathways, suggestive of an inter-dependent assembly of interacting proteins required for balanced signal branching. Given the similarity of LAT to other adapter-effector signaling systems, this mechanism may be a general paradigm to constrain signaling outputs in response to extracellular cues. Collectively, these insights extend our understanding of TCR-proximal signaling and present a basis for targeting LAT and similar proteins to tune T cell behavior in cancer, vaccination, and engineered cell therapies.

## Methods

### Generation of LAT knockout Jurkat T cell line

Jurkat Clone E6-1 cells (Jurkats) were purchased from ATCC and cultured in RPMI 1640 (Thermo Fisher 11875093) with 1x Pen/Strep/Glutamine (Thermo Fisher 10378016) and 10% FBS (Life Technologies 10082147). The Cas9-RNP nucleofection protocol from IDT was followed using a separate gRNA and tracrRNA, as well as Cas9 V3 products from IDT. The gRNA was designed using the Broad Institute sgRNA Designer tool and selecting the top hit (TTTACCAGTTTGTATCCAAG) which was predicted to create indels in Lat exon 4. The Cas9 RNP complex was generated following IDT’s recommendations and transfected into Jurkats using the Neon Electroporation System (Invitrogen). To isolate clones, cells were sorted into wells of a 96 round-bottom plate and left undisturbed in culture for two weeks. Wells with visible growth were resuspended, transferred to fresh media, and expanded. Thirteen clones were subjected to gDNA isolation and Illumina Nextera library preparation to sequence the locus surrounding the expected indel site. Primers LAT_sg2_Nextera_fwd and LAT_sg2_Nextera_rev (**Supplementary Table 2**) were used to perform PCR as a first stage, followed by an index PCR using Nextera P5 and P7 primers to add Illumina library and index sequences. Libraries were quantified by Qubit, pooled, and sequenced on a MiSeq (single end 130 base read length). The most abundant unique reads were aligned to the LAT locus using Benchling.

### Design and synthesis of cDNA library and expression vector

ORFs encoding the alanine block scan, targeted point mutations, GFP, charge alterations, and Cosmic/ClinVar human genetic variants were designed to be expressed in a modified version of the vector p_sc_eVIP (Addgene 168174)^1^. The 3’ UTR ORF barcode sequence was flanked by ATAC primer binding sites to enable the Spear-ATAC protocol, and a nested primary binding site was added downstream of the barcode to enable a nested PCR in both Spear-ATAC and CITE-seq ORF barcode library generation (**Figure S1B**). Twist Biosciences generated the individual expression vectors as their Clonal Genes product and shipped approximately 2-10 ug of each plasmid in separate wells.

### Preparation of titred lentivirus pool and transduction of Jurkat KO cells

HEK 293T (HEK) cells were cultured in DMEM (Thermo 11995065) with 10% FBS (Thermo Fisher 10082147) and 1x Pen/Strep/Glutamine (Thermo Fisher 10378016). Lentiviral packaging plasmids pMD2.G and pVSV-G were acquired from the Broad Institute Genetic Perturbation Platform. HEK cells were transfected using Lipofectamine 3000 (Thermo Fisher L3000001) following manufacturer’s protocol with modifications to achieve a 96-well format. To improve cell adherence, plates were coated with poly-L-lysine (Sigma P8920) for 1 hour at 37C (0.01% in PBS) followed by three washes with PBS and drying for 30 min at 37C. HEK cells were seeded at a density of 5.75e4 cells per well. The following day, cells were transduced by preparing lentivirus in a separate 96 well plate, first generating a standard packaging vector mix (10ul Optimem, 0.33ul P3000 reagent, 63ng pVSV-G, 21ng pMD2.G) and then depositing 83 ng of transfer vector (encoding each ORF) at 10 ng/ul to ensure consistent transfer to each well. A second mix was then deposited into each well containing 10ul Optimem and 0.42ul Lipofectamine 3000 and mixed gently by pipetting 7x. Manipulations were performed using a multi-channel pipet. Complex formation was allowed for 20 minutes at RT, and then 20ul of freshly prepared lipid complexes were added directly to culture media.

Plates were then cultured for 24 hours for a 48-hour post-transfection harvest, media was replaced, and a final harvest was taken at 72 hours post-transfection. An aliquot of the harvest was taken to perform p24 ELISA (Takara 631476) following manufacturer’s instructions on lentiviral supernatant diluted approximately 800-fold. ELISA values were computed to establish a mixing ratio for equal representation of viral particles across all library ORFs, and the ORF-specific lentiviral volumes were pooled for equal representation. The pool was then mixed with Lenti-X concentrator (Takara) following manufacturer’s instructions, concentrated, resuspended in PBS, and frozen in aliquots at −20C or −80C.

LAT knockout clone Jurkat cells (“Jurkat KO”) were transduced by mixing 1/24^th^ of the concentrated total viral harvest virus and 5e5 cells in media with 8 ug/ml polybrene. After overnight incubation, we performed a media change. At 48 hours post-transduction, cells were resuspended in media containing 0.6 ug/ml puromycin. Cells were cultured for approximately 7-9 days with passaging every 2-3 days to yield a population of viable (>90% by Trypan staining) cells. This amount of virus and transduction format led to roughly 10% transduction efficiency (as measured by proportion of viable cells at 24-48hr post-selection), suggesting most cells represent a single transduction event.

### TCR stimulation, CITE-seq and Spear-ATAC, and library generation

Jurkat KO cells transduced with the lentiviral ORF library pool were harvested and resuspended at 1e6 cells/ml in 1ml in a 12-well tissue culture plate to equilibrate for 30-60 minutes. Cells were then stimulated by adding anti-CD3 (OKT3, Thermo Fisher 16-0037-81) and anti-IgG (Biolegend 405301) antibodies at 1 ug/ml for 30 or 90 minutes before harvesting on ice. Cells were split into separate pools for CITE-seq and Spear-ATAC protocols. For CITE-seq, we followed the BioLegend protocol for simultaneous hashing and antibody staining (https://www.biolegend.com/en-us/protocols/totalseq-a-antibodies-and-cell-hashing-with-10x-single-cell-3-reagent-kit-v3-3-1-protocol). In brief, cells were blocked and stained with antibodies in Cell Staining Buffer (BioLegend 420201) and TruStain FcX (BioLegend 101319) using 1 ul of each antibody in 100ul for 30 min followed by three washes in Cell Staining Buffer (**Supplementary Table 2).**

Cells were then counted and cells from the 30-minute and 90-minute stimulation time points were pooled equally to allow for later distinction by hashing. Cells were loaded onto four channels of a 10x Genomics Chromium Next GEM Chip G at 3e4 cells per channel to leverage hash-based identification of doublets with super-loading. For Spear-ATAC, cells underwent nuclei isolation following the 10x Genomics suggested protocol for cell lines. Nuclei were counted and loaded on a 10x Genomics Chromium Next GEM Chip H at a concentration of 3e4 nuclei per channel (two channels per time point) to leverage identification of doublets by ORF barcode with super-loading. Following the Spear-ATAC protocol, addition of primer ORF-BC_nested_1_rev (**Supplementary Table 2**) was included in the encapsulation step and the number of stage one amplification cycles was extended to 15. Hashing (HTO), antibody (ADT), gene expression, and ATAC libraries were generated following 10x Genomics Chromium Next GEM Single Cell 3’ v3.1, BioLegend, and 10x Genomics Chromium Next GEM Single Cell ATAC v1.1 workflows.

### ORF barcode library generation from CITE-seq

Note: See **Supplementary Table 2** for primer sequences. NEBNext High-Fidelity 2X PCR Master Mix (NEBNext MM, New England BioLabs M0541S) was used for all PCRs. Generation of ORF identifying libraries from the CITE-seq experiment began by performing targeted amplification of the ORF construct from the cDNA material of the 10x Chromium Next GEM Single Cell 3’ v3.1 workflow (Step 2.2).

The first PCR was performed with the following reaction conditions:

Primers: CropDialOut_R1, BC_nested1_rev

Mix: 30ul reaction - 15ul NEBNext MM, 10uM each primer, 50ng cDNA product

Cycling: 98C for 30s; 6 cycles of 98C for 10s, 63C for 15s, 72C for 20s; 72C1min

This product was purified using 1x SPRIselect beads (Beckman Coulter B23317), with a 15ul elution in water, and used as input for a second nested PCR with the following reaction conditions:

Primers: CropDialOut_R1, BC_nested_Truseq_R2

Mix: 30ul reaction - 15ul NEBNext MM, 10uM each primer, 9ul PCR1 product

Cycling: 98C for 30s; 6 cycles of 98C for 10s, 63C for 15s, 72C for 20s; 72C 1min

This product was purified using 1x SPRIselect beads, with a 15ul elution in water, and used as input for a third indexing PCR with the following reaction conditions:

Primers: CropDialOut_P5_R1, P7_Truseq_idx[n]

Mix: 30ul reaction - 15ul NEBNext MM, 10uM each primer, 9ul PCR2 product

Cycling: 98C for 30s; 6 cycles of 98C for 10s, 63C for 15s, 72C for 20s; 72C 1min

This product was purified using 1x SPRIselect beads, with a 20ul elution in water.

These libraries were quantified by Qubit and Bioanalyzer (to identify the expected 379bp product) before mixing with CITE-seq gene expression, HTO, and ADT libraries for sequencing on a single Illumina NextSeq 2K P3 kit. The following reads per cell were targeted: gene expression – 1.8e3, ORF barcode – 1e3, ADT – 2.5e3, HTO – 1e3.

### ORF barcode library generation from Spear-ATAC

Note: See **Supplementary Table 2** for primer sequences. Generation of ORF identifying libraries from the Spear-ATAC experiment followed the published protocol with modification to the intermediate library purification strategy. Using the scATAC library as input, we performed a PCR using with reaction conditions:

Primers: P5_fwd, BC_nested_Truseq_R2

Mix: 30ul reaction - 15ul NEBNext MM, 10uM each primer, 50ng scATAC library

Cycling: 98C for 30s; 10 cycles of 98C for 10s, 63C for 15s, 72C for 20s; 72C 1min

This product was separated on a 2% TAE-agarose gel for one hour at 120V. A size range of 100 to 170 bp was excised (to capture the expected 119bp amplified fragment and exclude off-target amplicons), purified with a 22ul water elution, and used as input for a nested PCR with reaction conditions:

Primers: P5_fwd, P7_Truseq_idx[n]

Mix: 50ul reaction – 25ul NEBNext MM, 10uM each primer, 15ul purified PCR1

Cycling: 98C for 30s; 7 cycles of 98C for 10s, 63C for 15s, 72C for 20s; 72C 1min

This product was then purified using 2x SPRI beads and a 22ul elution in water. These libraries were quantified with Qubit and Bioanalyzer (to identify the expected 158 bp product) and sequenced on an Illumina MiSeq using a 150-cycle Reagent Kit v3. Custom primer CustomR1_PBS2 was used for read 1 (25 cycles) and ORF barcode information was extracted from read 2 (32 cycles), targeting roughly 1e3 reads per cell.

### FLAG-targeted inCITE-seq library generation

Jurkat KO cells transduced with the lentiviral ORF library (as described above) were harvested from culture and 1e6 cells were placed on ice. All subsequent steps (until loading onto the 10x Genomics chip) were performed on ice with pre-chilled reagents. PBS with FBS and recombinant RNase inhibitor (PBS/FBS/RRI) was prepared as the following mix: 8ml PBS, 160ul PBS (2% final), and 80ul SUPERase-IN RNase Inhibitor (Thermo AM2696, 0.2 U/ul final). Cells were spun down at 350g for 5 min, resuspended in 100ul fix buffer (497ul PBS/FBS/RRI, 3.13ul 16% PFA), and incubated on ice for 10 min. Then 300ul perm buffer (4.95ml PBS/FBS/RRI, 50ul 10% Tween) was added and cells were spun down at 200g for 5 min. Cells were then gently resuspended in 300ul perm buffer, incubated on ice for 5 min, and spun down at 200g. Cells were then gently resuspended in 50ul block buffer (249ul perm buffer, 1.25ul HCR probe hybridization buffer from Molecular Instruments). Anti-FLAG Total-seq A antibody (**Supplementary Table 2**) was added (0.5ul) and cells were stained for 20 minutes before two washes in 300ul perm buffer with 200g spins and gentle resuspensions. Cells were finally gently resuspended in 150ul PBS/FBS/RRI, filtered with a 100um FACS strainer, counted, and loaded on a 10x Chromium Next GEM Chip G with 3e4 cells per channel following the 10x Genomics Chromium Next GEM Single Cell 3’ v3.1 protocol. Generation of ADT and ORF barcode libraries was performed as described above.

### Bulk ATAC-seq

The Omni-ATAC protocol was followed largely as published protocol^2^. In brief, 5e4 cells from unstimulated or stimulated LAT KO Jurkat T cells transduced with LAT WT (as described above) were harvested on ice and subjected to nuclei isolation following the Omni-ATAC protocol. Purified ATAC fragments were amplified for eight cycles and purified using SPRIselect beads (1.2x). Libraries were quantified by Qubit and Bioanalyzer before sequencing on an Illumina NextSeq 500 targeting 2e6 read-pairs per library. The ENCODE ATAC-seq pipeline (https://github.com/ENCODE-DCC/atac-seq-pipeline) was used for pre-procesing, and chromVAR was used to infer transcription factor activities.

### Bulk RNA-seq

A modified version of the Smart-seq2 single-cell RNA-seq protocol was used^3^. Approximately 1e6 cells were harvested and total RNA was isolated using the RNeasy Mini Kit (Qiagen 74104). 10ng of RNA was used as input for the reverse transcription reaction and 12 cycles were performed for the whole transcriptome amplification step. The resulting cDNA (0.375ng) was used as input to the tagmentation reaction, followed by 12 cycles of indexing PCR. Libraries were purified using SPRIselect beads (0.9x), quantified by Qubit and Bioanalzyer, and sequenced on an Illumina NextSeq 500 targeting 2e7 read-pairs per library. The Cumulus Smart-seq2 pre-processing pipeline was used to generate gene expression count matrices which were depth normalized, centered, and scaled across samples for each gene^4^.

### Primary T cell isolation, genetic manipulation, and stimulation for single-cell analysis

Human primary CD4+ T cells were isolated from whole blood acquired from the Massachusetts General Hospital Blood Transfusion Service, following genomic data sharing policy guidelines and in accordance with the Broad Institute Office of Research Subject Protection (ORSP) protocol 3439. Peripheral blood mononuclear cells (PBMCs) were first isolated from blood using Ficoll-Paque PLUS density gradient media (Cytiva) following manufacturer’s instructions. CD4+ T cells were then isolated using the EasySep Human CD4+ T cell isolation kit (StemCell Technologies). Cells were either cultured directly in RPMI with 10% FBS, Pen/Strep/Glutamine (referred to as RPMI/FBS/PSG, as described above for Jurkat T cells) supplemented with 50 U/ml of IL2 (StemCell Technologies), or cryopreserved in BamBanker Freezing Media (Bulldog Bio). Prior to transduction, CD4+ T cells were expanded with Dynabeads Human T-Activator CD3/CD8 beads (Gibco) at a 1:1 bead:cell ratio, with approximately 1e6 cells in 1ml of media in a 24-well plate.

For transduction of primary T cells, lentivirus was generated in a similar fashion as described above for Jurkat T cells with several modifications. 4.5e6 HEK 293T cells were seeded in a 10cm dish (coated with poly-L-lysine as described above). The following day, cells were transduced with the Lipofectamine 3000 reagent (Thermo Fisher). For each sample, a first mix of 40ul Lipofectamine 3000 and 1ml Optimem media was made. A second mix of plasmids (8ug transfer plasmid, 6ug psPAX2, and 2ug pMD2.G (VSV-G) was diluted in 1ml Optimem, then 32ul P3000 reagent was added. The plasmid/P3000/Optimem mix was then pipetted on top of the Lipofectamine 3000 plus Optimem mix and incubated for 15 minutes at RT before the entire resulting volume was gently pipetted onto the cells. The following morning the media was changed. At 48hr and 72hr post-transfection, the supernatant was harvested, filtered with a 0.45 um low protein-binding syringe filter (Pall Corporation 4614), mixed with 3ml Lenti-X concentrator, and incubated at 4C for one to five days. The resulting mix was spun at 4C at 1500g for 45 minutes before supernatant was removed and the combined harvests from 48hr and 72hr were resuspended in 200ul RPMI/PBS/FBS with 50 U/ml IL2. These volumes were divided into 40ul aliquots and frozen at −80C.

Cells were transduced in a 96-well flat-bottom plate, allocating 10 wells for each virus (each encoding one ORF). Cells expanded with Dynabeads (1e6 cells in 1ml media) were first mixed by pipet to separate beads from cells. Then in each well we mixed 35ul cells (at approximately 1e6 cells per ml, not accounting for expected cell expansion after), 1ul Lentiboost (Sirion Biotech), 1ul thawed virus, and 49ul media with 50 U/ml IL2. This plate was spun for 1 hour at 32C at 931g and then incubated at 37C overnight. The following day, 100ul fresh media was added to each well and wells were pooled for each virus (2ml total) into a well of a 6-well plate. Dynabeads were then added (4e5) to achieve an approximate 2:1 ratio of beads: cells. After two more days of incubation, all cells were pooled, de-beaded using a magnet and counted. The resulting pool was divided into aliquots of 5e6 cells for nucleofection.

For nucleofection to knock out endogenous LAT using the Lonza 4D-Nucleofector, we assembled the ribonucleoprotein (RNP) complex using following a protocol similar that used to generate the Jurkat LAT KO cell line. We first annealed 1.65 ul crRNA (200uM) and 1.65 ul tracrRNA (200uM) with 4.2ul IDTE buffer to generate the guide RNA. This mix was heated at 95C for 5 minutes, then cooled to 75C at 1 C per second, then spun down and left at room temperature (RT) for five minutes. Cas9 V3 product from IDT was diluted in PBS at a ratio of 1.5:1 of Cas9:PBS. The RNP was then assembled by mixing equal volumes of annealed guide RNA with diluted Cas9 protein and incubating at room temperature for 10 to 45 minutes. IDT enhancer oligo was also diluted in P3 buffer (Lonza) by mixing 2.2ul enhancer oligo with 7.8ul P3 buffer. At this point, 3e6 cells were nucleofection sample were spun down in a 1.5ml tube and left at RT. Cells, RNP, fresh RPMI/FBS/PSG, a PCR strip tube, and the 16-well Lonza Nucleocuvette were brough to the nucleofection device. The nucleofector was set to cuvette mode with cell type specified as stimulated primary T cells, program EH-115, and buffer P3. For each sample, 1ul of enhancer oligo was mixed with 1ul of RNP in a strip tube. Then one cell pellet sample was gently resuspended in 20ul P3 buffer and that volume was transferred to the strip tube containing enhancer oligo and RNP. The total resulting volume was mixed gently and transferred to one well of the Nucleocuvette carefully to avoid bubbles. After loading all samples, the Nucleocuvette was placed in the nucleofector, the program was run, and the Nucleocuvette was quickly removed. 100ul RPMI/FBS/PSG (no IL2) was dripped into each well without mixing, and the Nucleocuvette was transferred to a 37C tissue culture incubator for 15 min. Next the cells were transferred to a well of a 12-well plate containing pre-equilibrated RPMI/FBS/PSG with 500 U/ml IL2. The following morning, cells were spun down and resuspended in 1ml RPMI/FBS/PSG with 50 U/ml IL2 and incubated overnight. The following day, all cells were pooled, spun down, and resuspended in RPMI/FBS/PSG with 50 U/ml IL2 to a concentration of 1.2e6 cells per ml.

After two more days, cells were spun down and resuspended at 1.5e6 cells per ml with 0.6 ug/ml puromycin to begin selection. After two more days, cells were counted (viability ∼50%). These cells were processed using the EasySep Dead Cell Removal Annexin V Kit (StemCell Technologies), which increased viability to ∼70% and cells were maintained at 1e6 cells per ml with puromycin. At this point a sample of cells was also taken to confirm Cas9 activity by genomic DNA extraction, PCR amplification of the targeted locus, and Sanger sequencing. Using TIDE^5^ to analyze Sanger sequencing results, we achieved ∼75% indel generation.

After two more days, cells were harvested and prepared for TCR stimulation followed by single-cell RNA-seq and ATAC-seq analysis on the 10x Genomics platform, similar to the workflow performed on Jurkat T cells describe above. 2.5-5e5 cells were resuspended in 500ul of media with 50 U/ml IL2 and rested for one to three hours. Then anti-CD3 and anti-IgG antibody were added to wells and mixed to achieve a final 1 ug/ml concentration of each antibody. Cells were incubated for 90 minutes before harvesting. For scRNA analysis, cells were stained with BioLegend Hashtag B antibodies with barcodes 1-4 (following the same protocol for CITE-seq antibody staining described above for Jurkat T cells) to identify each of four replicate samples.

Each replicate sample was counted, mixed evenly, and diluted to 1e3 cells per ul before loading 30ul of cells onto two channels of Chip G using the 10x Genomics scRNA v3.1 kit. Gene expression and ORF barcode libraries were generated as described above, and feature barcoding hashtag library was generated following 10x Genomics suggested protocol. For scATAC analysis, cells were processed in a similar fashion as the Jurkat T cell experiment described above. 1.8e4 cells from each of two replicate samples were loaded into two channels of Chip H (one sample per channel) using the 10x Genomics scATAC v2 kit. Further processing with Spear-ATAC modifications for ORF barcode recovery was performed as described above for the Jurkat T cell experiment.

### LAT protein proximity labeling with TurboID in activated T cells

The lentiviral expression vector described above was modified to express LAT with a C-terminal TurboID fusion. In brief, we designed an IDT gBlock Gene Fragment encoding a glycine-serine linker, TurboID (sequence derived from Addgene plasmid 107169), and a V5 tag (**Supplementary Table 2**). A PCR amplicon encoding LAT with overhangs for cloning was generated using primers Fwd_LAT_forTurboID and Rev_LAT_forTurboID. The amplicon was agarose gel purified, and the product was used for an In-Fusion (Takara Bio) cloning reaction with the following components: 1ul In-Fusion master mix, 1ul of transfer vector (digested with NheI for 2 hrs and gel purified) at 100 ng/ul, 0.5ul LAT PCR amplicon at 70 ng/ul, 0.5ul TurboID gBlock at 50 ng/ul, and 2 ul water. After a 15 minute incubation at 50C, 2.5ul of this reaction was used to transform approximately 20ul of NEB Stable Competent E. coli. Bacteria were grown at 30C to prevent recombination of lentiviral repeat regions. A single colony was picked from an LB agar plus ampicillin plate and grown overnight in LB media with ampicillin for miniprep and Sanger sequencing validation.

To detect LAT proximal proteins by western blot we followed an established protocol^6^ with some modifications. Lentivirus was generated using the LAT-TurboID transfer vector as described above, and LAT KO Jurkat T cells were transduced and selected as described above. For each stimulation and labeling experiment, 5e6 transduced cells were rested for 1 hour in 1ml in a 12-well plate in a 37C incubator. We used the phosphatase inhibitor pervanadate to stimulate cells, an approach which mimics direct TCR stimulation but yields a stronger, more durable response to facilitate protein interaction detection^7–9^.100x concentrated pervanadate was made as a mix of 300 mM hydrogen peroxide (from a fresh bottle within ten days of opening) and 10 mM sodium orthovanadate in water. We then added 10ul of 100x pervanadate to 1ml of cells, mixed the samples, and returned the plate to the incubator for ten minutes of labeling. No exogenous biotin was added to the culture (the level of biotin in RPMI 1640 media was sufficient for robust labeling). The plate was then placed on ice, and cells were transferred to 1.5ml tubes for three washes in 1ml of DPBS at 4C at 400g. For each wash, we took care to remove all supernatant by first removing 900ul of supernatant for all samples and then removing the remaining ∼100ul in a quick aspiration step. Cells were then lysed by resuspending in 100ul RIPA buffer with protease inhibitor cocktail (Millipore Sigma P8849) and PMSF (VWR 82021-256), thoroughly pipet mixing, and incubating on ice for 10 minutes. Samples were then spun at 4C at 13e3 g for 10 min to clarify the lysate. 5ul of clarified lysate was taken as an input sample and mixed with 5ul reducing Laemmli buffer (Boston Bioproducts BP-111R) and 20ul water, then boiled for 5 minutes at 95C. 90ul of clarified lysate was mixed with Pierce Streptavidin Magnetic Beads (Thermo Fisher 8816) resuspended in 500ul RIPA buffer (25ul beads stock per sample) in a 1.5ml tube. Streptavidin pull-down of biotinylated proteins was performed for two to three hours by rotating samples at 4C. Bead washing was performed using a magnetic rack in an ice bucket with buffers pre-chilled and 900ul buffer per wash. For each wash, tubes were quickly transferred from one side of the magnet rack to the other and then returned to the original side to ensure bead mixing. The following washes were performed: two washes with RIPA (two transfers each), one wash with 1M KCl (two transfers), one wash with 0.1M Na2CO3 (one transfer), one wash with 2M urea (in 10mM Tris-HCl pH 8) (one transfer), and two washes in RIPA (two transfers each). After removal of the last wash buffer, tubes were quickly spun down with a bench-top centrifuge and returned to the magnet to remove all supernatant. Then 40ul elution buffer (150ul 6x Laemmli buffer, 6ul 0.1M biotin, 144ul water) was added to each well and samples were spun again quickly. A p20 pipet was then used to thoroughly resuspend the beads in elution buffer, with vigorous pipetting approximately ten times. Samples were then boiled for 10 min at 95C to elute proteins. Tubes were spun again quickly and returned to the magnet, and after 30 seconds the elution volume was transferred to a fresh tube.

We then ran a western blot using the NuPAGE Bis-Tris gel system (Thermo Fisher) with 5ul input and 15ul enrichment for each sample along with 4ul Precision Plus Protein standard (Bio-Rad 1610374). The gel was separated at 180 V for 30 minutes, followed by a transfer using the iBlot2 system nitrocellulose membranes with the P0 protocol. The membrane was cut horizontally at pre-identified sizes for PLCG1 (Cell Signaling 2822S), GAPDH (Thermo Fisher PA1-987), and GRB2 (Thermo Fisher PA5-27151), and incubated overnight with 1:1000 primary antibody in TBS-T with 5% milk at 4C on a rocking mixer. The following day, membranes were washed three times in TBS-T for 5 minutes each and incubated in 1:2000 secondary antibody targeting Rabbit IgG conjugated to HRP (Thermo Fisher A27036) in TBS-T with 5% milk at RT on a rocking mixer. Membranes were again washed three times in TBS-T for 5 minutes each, and ECL reagent (Sigma GERPN2232) was used to visualize proteins on a Bio-Rad ChemiDoc MP imager. Bands were quantified using the FIJI Analyze Gels function.

Normalized enrichment values were calculated by dividing the enrichment band value by the mean of the input value for GAPDH and PLCG1. Means and error bars in Figure 4 represent values from four replicate experiments performed on four separate days.

### Jurkat T cell TCR stimulation and flow cytometry validation of mutant phenotypes

LAT KO Jurkat T cells were transduced with lentivirus expressing a single ORF and selected with puromycin as described above. 2e5 cells per sample were deposited in two replicate wells for an unstimulated condition and two replicate wells for a stimulated condition in 200ul media in a 96-well flat-bottom plate. Cells were equilibrated for three hours before addition of anti-CD3 antibody for a final concentration of 0.1 ug/ml and mixed with a multi-channel pipet. After six hours, cells were transferred to a v-bottom 96-well plate, spun at 350g for 5 min at 4C in a swinging-bucket centrifuge, and resuspended in 50ul PBS plus 2% FBS containing 1:100 diluted anti-CD69-APC antibody (BioLegend 310909). Cells were stained in the dark on ice for 30 min, then 200ul PBS + 2% FBS was added and cells were washed twice in the same buffer before a final resuspenion in 200ul of the same buffer. Cells were analyzed on a Beckman CytoFLEX flow cytometer. Mean APC values were calculated for each sample and normalized to value for cells expressing wild type LAT.

### Single-cell RNA-seq and ATAC-seq data pre-processing and ORF barcode assignment

For gene expression, ADT, HTO, and scATAC libraries, Cellranger pre-processing with standard parameters from the Cumulus V2.1.1 instance was used to generate cell-level expression matrices as well as chromatin accessibility peaks and fragment files^4^. Seurat v4.3.0 was used to process scRNA, ADT, and HTO libraries^10^. Cells were filtered for number of genes (between 5e2 and 1e4) and mitochondrial content (up to 10%). Gene expression matrices were then normalized using SCTransform in Seurat using parameters vst.flavor = “v2”. SCT residuals were used for all further analyses. Signac v1.9.0 was used with Ensembl database EnsDb.Hsapiens.v86 and hg38 to process scATAC data^11^. Cells were filtered for peak region fragments (1e3 to 3e4), percent reads in peaks (>15%), nucleosome signal (<4), and TSS enrichment (>2). ChromVAR was run using the JASPAR 2020 motif database and deviation values were used as inferred TF (motif) activities for all further analyses.

Custom UNIX and Python scripts were used to process fastq files from ORF barcode libraries into per-cell counts of ORF barcodes from a register of the library barcodes. ORF barcode counts were normalized by total counts per cell, and cells were thresholded on a minimum total counts for CITE-seq (50 reads) and proportion of reads supporting the top ORF barcode (80% for Spear-ATAC and CITE-seq) based on analysis of the distribution of proportions. The top ORF barcode in each cell was assigned as the ORF identity.

### Generation and statistical analysis of ORF-level activation scores and feature values

A single activation score for each cell was calculated for each time point. To generate a chromatin score from ATAC data, the top 50 TF motifs with greater chromVAR signal in cells expressing wild type LAT versus cells expressing GFP were averaged. For RNA samples, the top 50 genes with greater SCT residuals in cells expressing wild type LAT versus cells expressing GFP were averaged. To determine statistical significance of scores deviating from wild type LAT at the level of ORFs, the scores for each cell expressing an ORF were averaged and 1000 samples, matched to the number of cells supporting the ORF being examined, were taken from each of the three wild type LAT replicate cell pools. For each sample, the average score across cells was calculated and the difference in score between sampled cells and the WT replicate pool from which it was sampled was determined. For each WT replicate sample, we counted the number of instances in which the sample had a more extreme value than the value observed for the cells expressing the ORF, and this proportion was recorded as the false discovery rate (FDR). The same approach was used to determine FDRs for individual feature values (TF motifs, genes, or CD69 protein).

To determine ORF clusters, a matrix of chromatin and RNA scores from each time point was clustered using the R k.means function with default parameters and k=3.

### Annotation of LAT protein sequence features with the Eukaryotic Linear Motif Database and localCIDER

The amino acid sequence of LAT (ENST00000395456) was entered into the Eukaryotic Linear Motif resource website (www.elm.eu.org) using all cell compartments and a default motif probability cutoff of 100. The resulting table was downloaded and processed in R to display the visualization in Figure 2. LAT homolog protein sequences were extracted from the ConSurf tool output (https://consurf.tau.ac.il/consurf_index.php). To determine amino acid sequence identity of each homolog compared to human LAT, NCBI BLASTP was performed using default parameters. A random set of human length-matched proteins was derived from the UniProt database website after filtering for homo sapiens (organism ID 9606) and “Reviewed” proteins. The resulting fasta file was processed in Python to extract 100 random proteins with length between 200 and 250 amino acids to approximate the length of LAT. To calculate biophysical sequence features, localCIDER was run using Python to analyze all LAT homolog sequences and random background proteins.

### Prediction of protein binding to LAT with AlphaFold-Multimer

AlphaFold-Multimer was run using Colabfold V1.5.5 via Docker (ghcr.io/sokrypton/colabfold:1.5.5-cuda11.8.0) on a Google cloud project virtual machine (NVIDIA T4 GPU with 30GB memory)^12^. We generally followed instructions described on the associated GitHub (https://github.com/sokrypton/ColabFold/wiki/Running-ColabFold-in-Docker). Predictions were generated without templates or relaxation, and MMseqs2 server was used for multiple sequence alignments (paired and unpaired). Sequences for full length proteins and SH2 domains were retrieved from Uniprot. To further interpret the output of Colabfold, we implemented a recently described workflow implemented in Python which parses the structure files (PBD format) and AlphaFold-Multimer confidence metrics to identify confident contact regions between two protein chains^13,14^. To generate ten unique models (instead of the maximum of five) when analyzing isolated SH2 domains interacting with LAT, we ran Colabfold a second time with a different random seed. The 10 PDB files representing 10 output models from each SH2 domain were then analyzed using a custom Python script in Pymol to determine the distance between each LAT tyrosine and residue 48 in the SH2 domain, which consistently represented the proximal residue to the LAT tyrosine in interacting structures. A structure was scored as an interaction if the SH2 domain distance to the most proximal tyrosine was less than 10 angstroms and the second nearest tyrosine was at a distance greater than ten angstroms.

### Determination of coordinated and biased signaling outputs from chromatin accessibility data

To determine whether individual ORFs conferred coordinated (loss of both AP-1 and NFAT TF activity) or biased (loss of only one TF activity), inferred TF activity was first scaled similarly to the chromatin activation score, such that 0 represented the mean value of GFP-expressing cells and 1 represented the mean value of WT LAT-expressing cells. These values were then averaged across the 30 and 90 minute stimulation experiments, and only ORFs supported by more than 25 cells in each time point were analyzed. An ORF was considered to confer a significant defect for a TF activity if the permutation-based FDR (described above) met a threshold of 0.05 in both time points. ORFs were classified as coordinated if AP-1 (FOS motif) and NFAT (NFATC1 motif) were significantly defective. An ORF was classified as biased if either AP-1 or NFAT were significantly defective in both time points and the other TF had a confidently unaltered response with an FDR threshold of 0.1 in both time points.

### Figure generation

Figures were generated using Adobe Illustrator and BioRender.

## Acknowledgments

We thank members of the Shalek and Regev labs for helpful discussions. We thank J.T. Neal, Bingxu Liu, Matteo Gentili, and Nir Hacohen for sharing experimental resources and guidance. A.J.R was supported by the Helen Hay Whitney Foundation Research Fellowship. T.T.D. was supported by the United States Department of Agriculture Agricultural Research Service Graduate Student Fellowship. A.K.S was supported, in part, by the NIH NIDA DP1 Avant-Garde Pioneer Award (1DP1DA053731) and NIAID 1R01AI149670; the Bill and Melinda Gates Foundation; and the Ragon Institute of MGH, MIT, and Harvard. A.R. was supported by the Klarman Cell Observatory at the Broad Institute of MIT and Harvard.

## Author Contributions

A.J.R, A.R., and A.K.S. conceived the project. A.J.R, T.T.D, and A.V.S performed experiments and analyzed data. A.K.S. and A.R. guided experiments and data analysis. A.J.R, T.T.D, A.V.S., A.R., and A.K.S wrote the manuscript.

