## Supplementary Figures 1-4 for "LAT encodes T cell activation pathway balance"

### Supplementary Figure 1

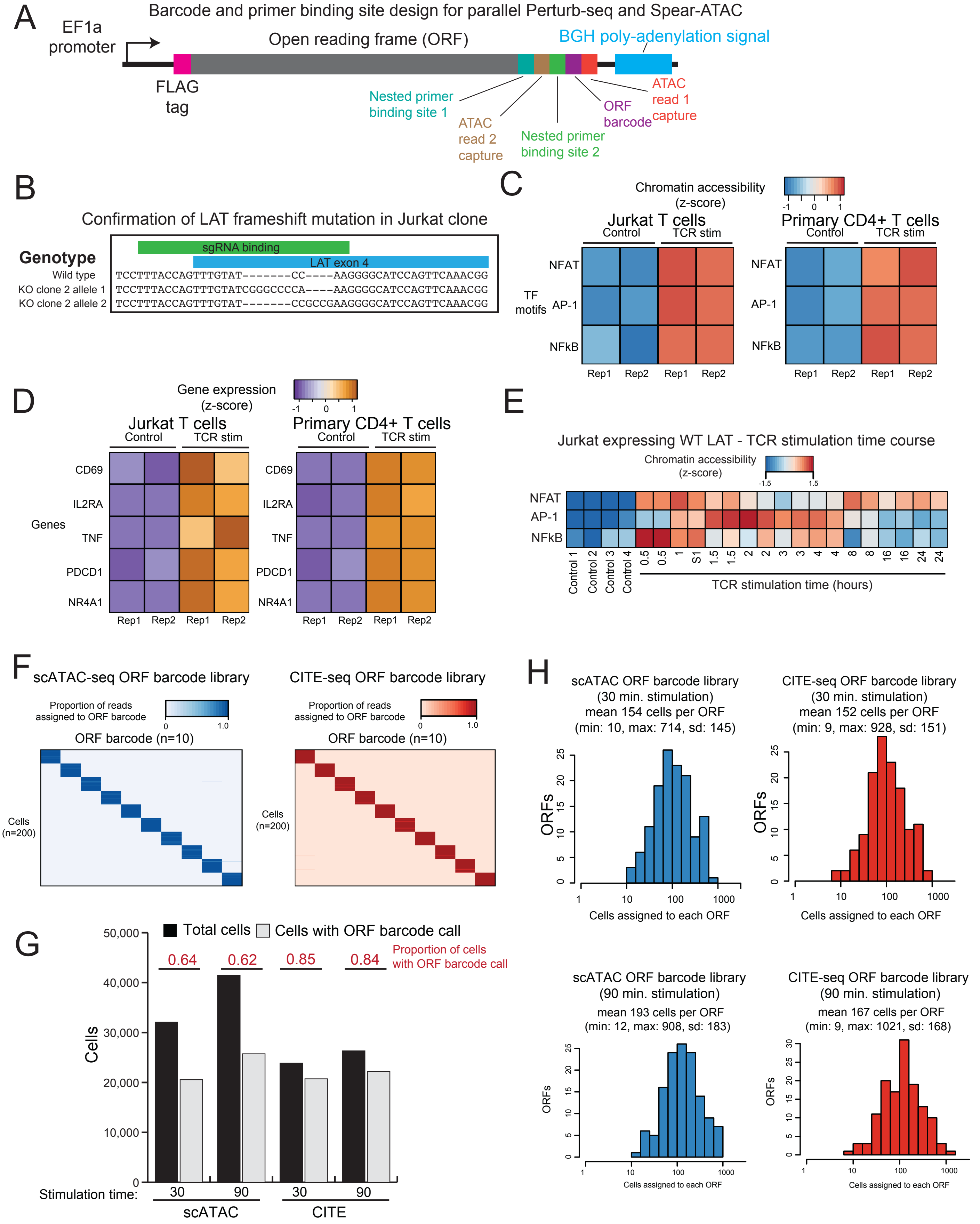

#### Supplementary Figure 1

(**A**) Schematic of ORF expression vector including a 3' UTR containing the ORF barcode and primer binding sites to amplify the ORF barcode in Spear-ATAC and CITE-seq assays. (**B**) Wild type DNA sequence of LAT with sequencing reads from top identified alleles in knockout clone 2 are aligned. sgRNA binding site is indicated, and each allele has an out of frame insertion. (**C**) Bulk chromatin accessibility analysis on Jurkat T cells (ATAC-seq) and primary human CD4+ T cells (DHS-seq from ENCODE project). Each row corresponds to a set of genomic regions with a shared TF motif. Chromatin accessibility quantified by chromVAR. TF motifs correspond to NFATC1 (NFAT), FOS (AP-1) and REL (NF-kB). (**D**) Bulk RNA-seq on Jurkat T cells and primary human CD4+ T cells. (**E**) Heatmap of relative chromVAR TF motif accessibility values generated from bulk ATAC-seq profiling of Jurkat T cells transduced with WT LAT. Cells were either left unstimulated (control) or stimulated for 0.5 to 24 hours. TF motifs correspond to NFATC1 (NFAT), FOS (AP-1) and REL (NFκB). (**F**) Detection of ORF barcodes in single cells from the Spear-ATAC (left) or CITE-seq (right) libraries. Rows represent subsampled cells from each experiment, columns represent a subset of ORFs in the library. Each cell has a high proportion of reads devoted to a single ORF barcode. (**G**) Bar plot indicating total cells passing quality filters and cells assigned an individual ORF. (**H**) Histograms of the number of cells assigned to each ORF in the 30 min. or 90 min. stimulation scATAC or CITE-seq libraries.

### Supplementary Figure 2

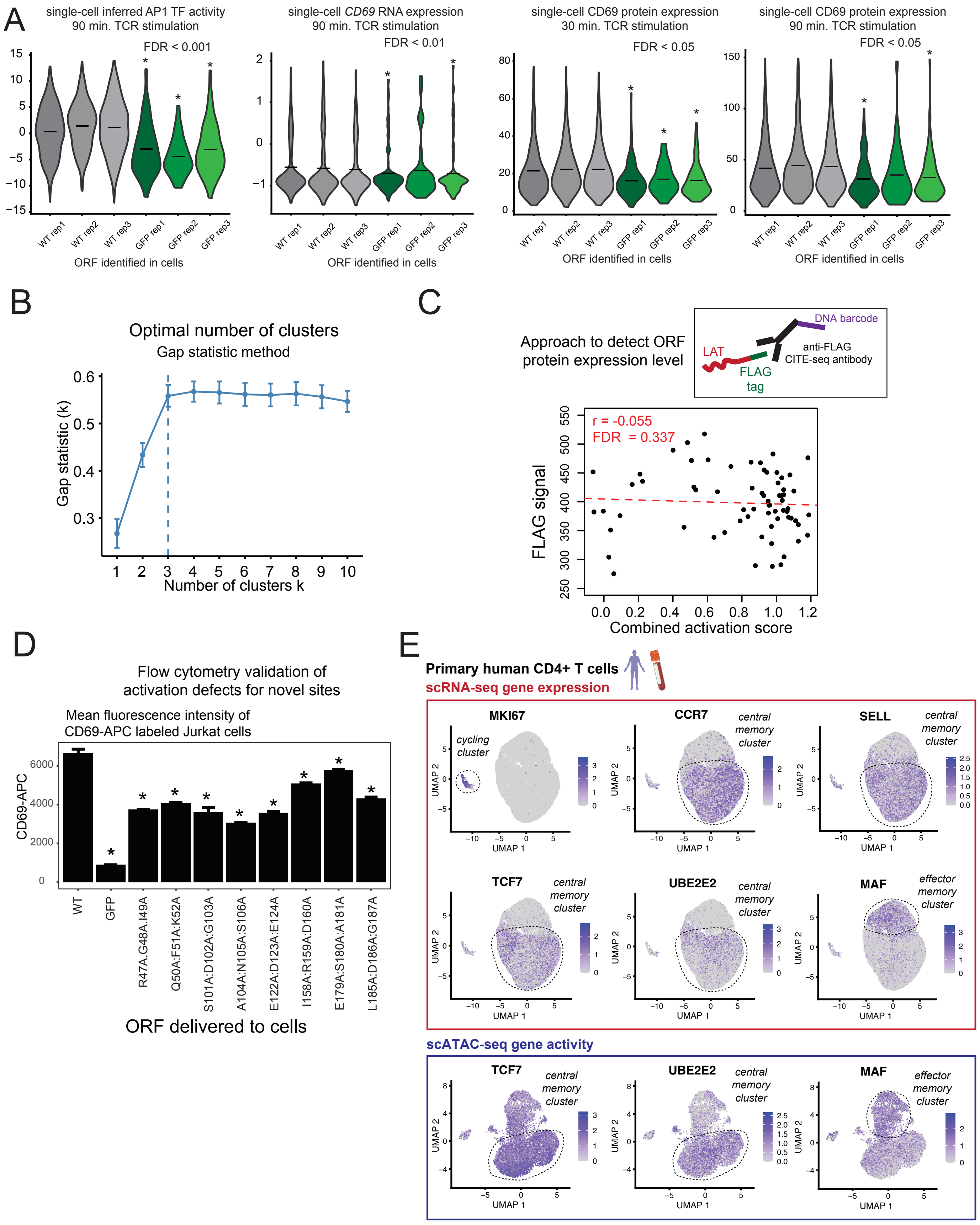

#### Supplementary Figure 2

(**A**) Similar to Figure 1D,E, violin plots representing single-cell inferred TF activity for AP-1, RNA gene expression for *CD69*, or surface protein expression of CD69. Line indicates mean across cells and FDR generated from single-cell sampling test. (**B**) Gap statistic values for identifying the optimal number of k-means clusters. Calculated using the NbClust and factoextra R packages. (**C**) Top: Schematic of inCITE-seq approach to monitor protein expression levels of all ORFs. Bottom: Scatter plot of combined activation score versus protein expression detected by anti-FLAG tag targeted inCITE-seq. FDR calculated by permuting ORF identities indicates ORF protein expression level is not correlated with activation defect. (**D**) Bar plot representing cell surface expression of CD69 measured by flow cytometry. LAT KO Jurkat cells were separately transduced with lentivirus expressing WT LAT, GFP, or the listed LAT mutants. Cells were stimulated with anti-CD3 antibody for six hours before staining with anti-CD69-APC antibody for flow cytometry. Values represent mean across two technical represents and error bars indicate standard deviation. *, $p < 0.05$ from two-tailed t-test. (**E**) UMAP dimensionality reduction of primary human CD4+ T cells from single-cell RNA-seq and ATAC-seq experiments. Cells color is scaled to indicate the expression level of individual genes in the scRNA-seq experiment or inferred gene activity from the scATAC-seq experiment. Groups of cells are circled to indicate group labels. Shared markers TCF7 and UBE2E2 between scRNA and scATAC experiments indicate central memory T cells, while shared marker MAF indicates effector memory T cells.

### Supplementary Figure 3

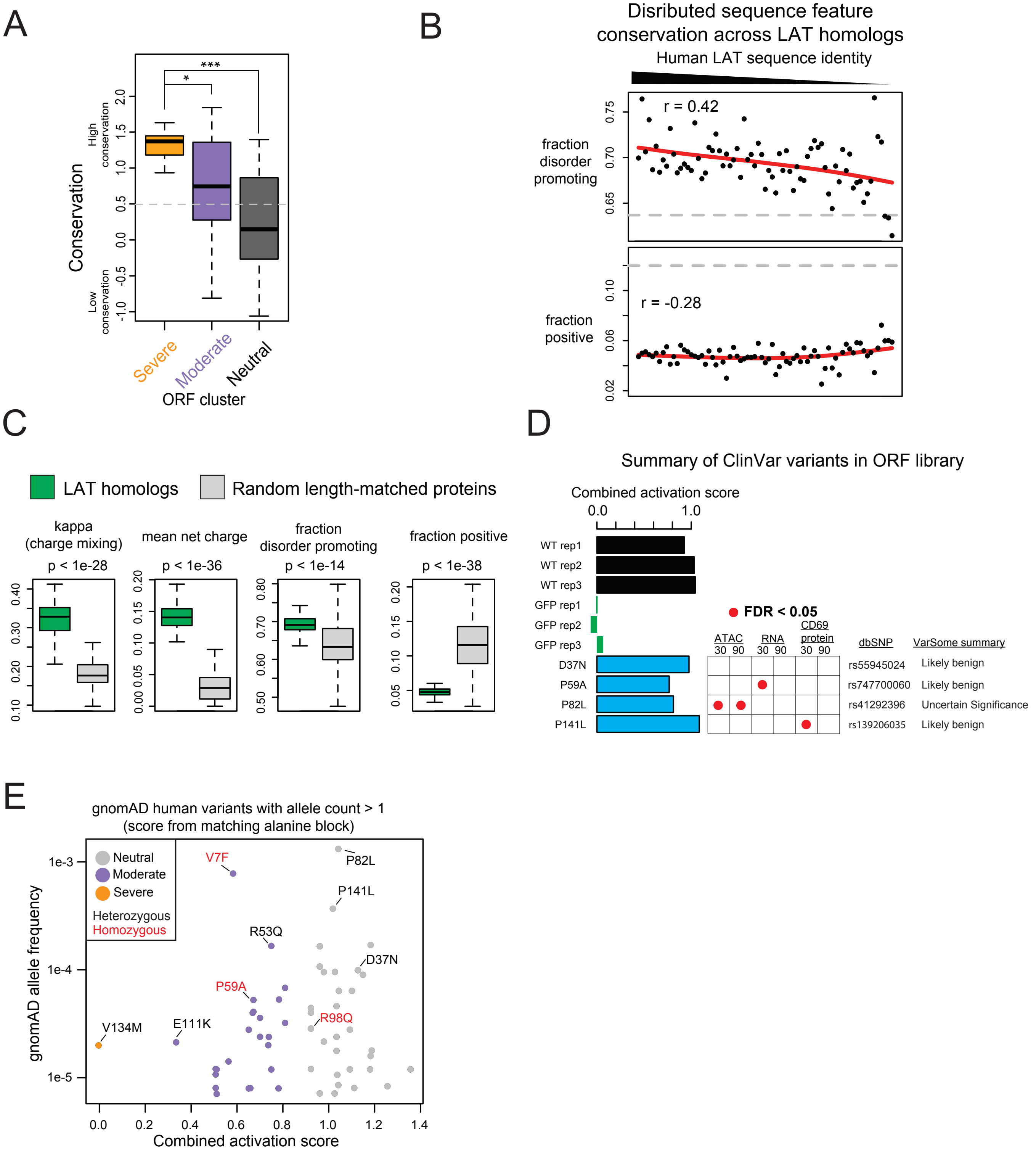

**Supplementary Figure 3**

(**A**) Box plot of ConSurf conservation scores for amino acids mutated in ORFs from each functional cluster. **, KS test p-val < 0.01; *, KS test p-val < 0.05. (**B**) Scatter plots of CIDER distributed amino acid sequence features for LAT homologs, ordered by amino acid sequence identity with human LAT.  Pearson correlation (r) calculated between sequence identity percent and each feature. Red line indicates a smooth spline calculated in R and gray line indicates mean of random human length-matched proteins. (**C**) Box plots of CIDER features comparing LAT homologs to random length-matched human proteins. P-values represent KS test. (**D**) Combined activation score of LAT WT, GFP, and ORFs encoding ClinVar missense variants. VarSome database high level prediction of function based on aggregation of prediction tools. (**E**) Scatter plot of combined activation score of alanine blocks and corresponding allele frequency for gnomAD database missense variants aligned to alanine blocks. Variants colored in red were found in individuals as homozygous at least once.

### Supplementary Figure 4

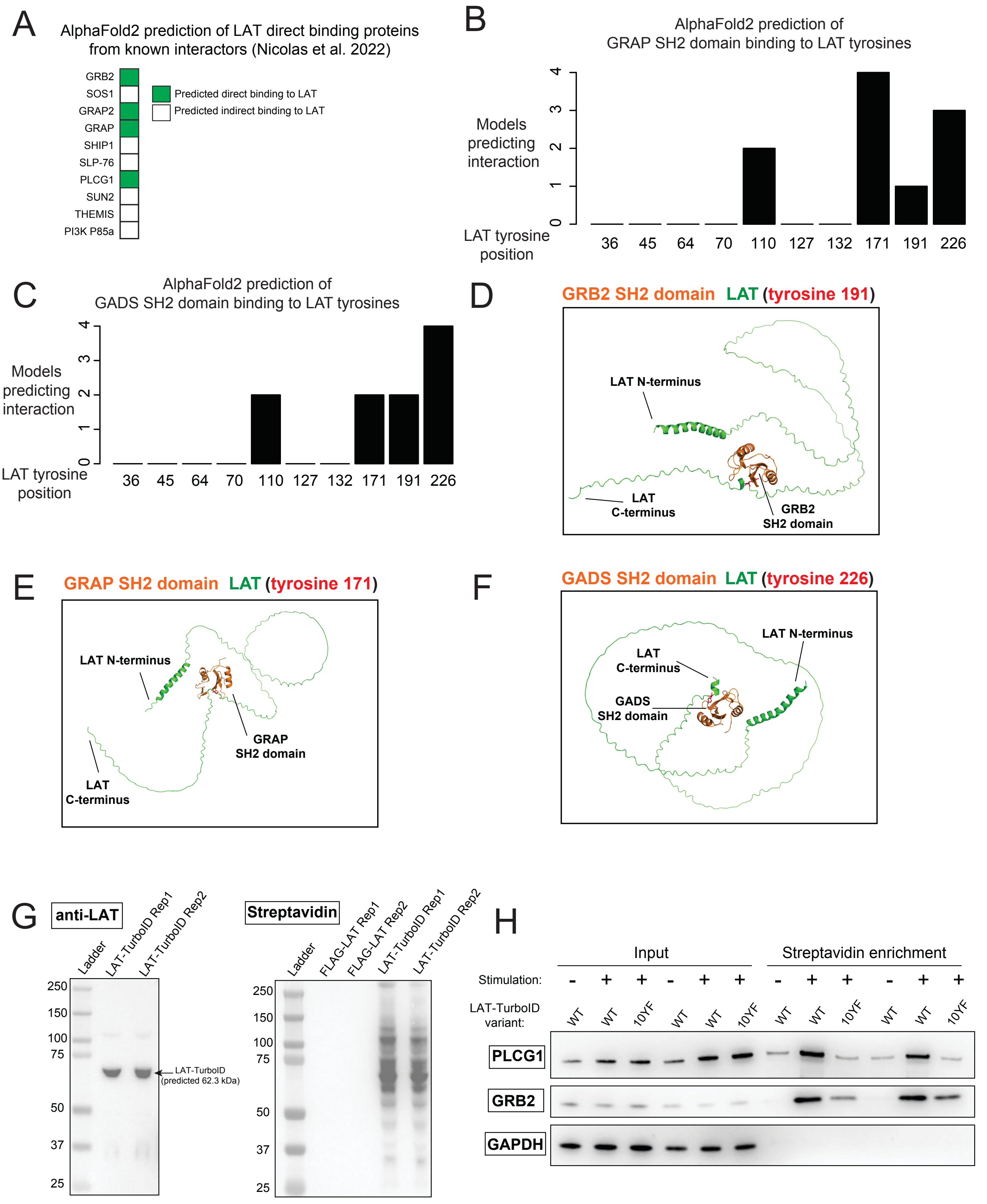

#### Supplementary Figure 4

(**A**) Summary of AlphaFold-Multimer survey of known LAT interacting proteins. All proteins found to interact with LAT via affinity purification mass spectrometry (from Nicolas et al. 2022) were used as input to AlphaFold-Multimer to predict a binary interaction structure with LAT. (**B**) Counts of models (out of 10 total predicted models) for each LAT interaction with the SH2 domain of GRAP. Models were scored as interaction with a particular LAT tyrosine based on exclusive proximity of 10 angstroms. (**C**) Similar to (B), for LAT interaction with the SH2 domain of GADS. (**D**) Predicted structure of LAT interacting with the GRB2 SH2 domain. (**F**) Predicted structure of LAT interacting with the GRAP SH2 domain. (**E**) Predicted structure of LAT interacting with the GADS SH2 domain. (**F**) Western blot using anti-LAT antibody to detect expected full-length expression of LAT-TurboID fusion protein. (**G**) Western blot using streptavidin-HRP to detect biotin labeling exclusively in cells expressing the LAT-TurboID fusion. (**H**) Western blots of expected LAT interacting proteins PLCG1 and GRB2, along with GAPDH, in LAT-TurboID labeling experiment. Cells were stimulated with pervanadate for 10 minutes before lysis and streptavidin enrichment to detect labeled proteins. 10YF indicates LAT mutant with all ten tyrosine positions mutated to phenylalanine, which is expected to disrupt all binding between LAT and partner proteins mediated by partner SH2 domains.
